# Proportional recovery in mice with cortical stroke

**DOI:** 10.1101/2024.10.11.614428

**Authors:** Aref Kalantari, Carolin Hambrock, Christian Grefkes, Gereon R. Fink, Markus Aswendt

**Author notes:** Correspondence: Markus Aswendt, PhD, University Hospital Cologne, Department of Neurology, Kerpener Strasse 62, 50937 Cologne, Germany.

## Abstract

The proportional Recovery Rule (PRR) has been frequently used to predict recovery of lost motor function in acute stroke patients. However, it still needs to be explored whether the same concept applies to preclinical, i.e. animal models of stroke recovery. To address this question, we investigated behavioral data from 125 adult male C57Bl/6J mice with photothrombotic strokes in the sensorimotor cortex. Lesion size and location were determined in the first week using in vivo T2-weighted MRI. Motor recovery was evaluated repeatedly over four weeks using the cylinder, grid walk, and rotating beam test. Recovery trajectories were analyzed using a newly formulated Mouse Recovery Rule (MRR), comparing it against the traditional PRR. Initial findings indicated variable recovery patterns, which were separated using a stepwise linear regression approach resulting in two clusters: 47% PRR and 53% MRR. No significant correlation was found between recovery patterns and lesion size or location, suggesting that other biological factors drive individual differences in recovery. Of note, in the MRR cluster, animals recovered to 90% of their initial behavioral state within the first four weeks post-stroke, which is higher than the 70% recovery usually reported in human PRR studies. This study demonstrates the complexity of translating the PRR to stroke recovery models in mice and underscores the need for species-specific recovery models. Our findings have implications for designing and interpreting therapeutic strategies for stroke recovery in preclinical settings, with the potential to improve the predictive accuracy of stroke recovery assessments.

## Introduction

The mechanisms driving recovery from focal brain lesions as in stroke are still incompletely understood despite intensive research over the past decades (Grefkes and Fink 2020). A major problem is that recovery profiles in preclinical models often differ from those observed in patient populations, thereby hampering interspecies comparisons and translation. An important and widely used concept in human stroke recovery research is the proportional recovery rule (PRR) which posits that the magnitude of recovery from motor impairment is approximately 70% of the initial impairment (Prabhakaran et al. 2008). However, applying the PRR across studies has yielded inconsistencies, necessitating methodological caution. A common approach in applying the PRR involves fitting a linear regression model that relates initial impairment to change in impairment. Concerns have been raised regarding the removal of ‘non-recoverers’ or ‘non-fitters’, regression diagnostics, and potential nonlinear associations between recovery and initial impairment (Bonkhoff et al. 2022; Bowman et al. 2021; Hope et al. 2019; Kundert et al. 2019). There is a mismatch between clinical studies supporting the generalizability of the PRR in larger cohorts (Kundert et al. 2019), and only a handful of preclinical studies in monkeys and rats (Jeffers, Karthikeyan, and Corbett 2018; Nashed et al. 2024).

Animal models, particularly rat and mouse models, are believed to be strongly limited in the application and interpretation of the PRR (Balkaya and Cho 2019). To our knowledge, there is only a single study of the PRR in a large cohort of n=593 male rats using a unilateral endothelin-1 lesion model and functional assessment using the Montoya staircase-reaching task (Jeffers et al. 2018). In contrast to human studies, a smaller part of approx. 30% were identified as fitters, characterized by smaller infarct volumes and initial post-stroke impairments compared to non-fitters. This finding suggested that the existing motor measures, unlike the FM-UE used in humans, might not reflect the actual impairment adequately. The term “recovery” in rodent studies often refers to improved performance on tasks that do not distinguish spontaneous – or treatment-induced recovery from compensation (Corbett et al. 2017). In contrast to the standardized Fugl-Meyer upper limb score (FM-UE) in humans, scoring systems in mice, such as the Bederson scale of forelimb flexion (Bederson et al. 1986), are somewhat subjective, less standardized, and not widely used. For example, compensatory responses, e.g., avoiding impaired limb use or using a different movement pattern, are hard to distinguish from “true” recovery (Jones 2017), especially when with single rodent motor tests focused on a specific movement (e.g. gait). Therefore, applying a battery of motor tests, including spontaneous limb movements (e.g. in the cylinder test), is recommended to detect compensatory mechanisms (Corbett et al. 2017).

To address the gap of PRR validation in a stroke mouse model, we transformed three sensorimotor tests, well established to assess motor recovery after cortical stroke, i.e., the cylinder, grid walk, and rotating beam test, applied over four weeks. We measured the ratio of mice following the PRR and determined a group with accelerated recovery in relation to stroke lesion size and location.

## Methods

### Experimental model

The experimental protocol was designed according to the IMPROVE and ARRIVE principles. Adult male C57Bl/6J mice (9-15 weeks old, The Jackson Laboratory) were randomly assigned to the stroke and sham groups. For this study, several experiments with the same stroke mouse model were pooled based on an initial pre-screening including only mice with longitudinal data (MRI and behavior) from at least three time points. See detailed description in **Supplementary Material**.

### Deficit Score Calculation

To accurately quantify the behavioral state of the mice, a single deficit score was introduced, which combines the results of the rotating beam, cylinder, and grid walk tests. Because these tests depict different types of sensorimotor deficits, which are analyzed differently and therefore have different scales, z-transformation was used to normalize the scores. This way it was possible to average the results of each mouse per test into one composite score, referred to as the deficit score.

### Application of the Proportional Recovery Rule

To describe mathematically the proportional recovery rule (PRR) based on the Fugl-Meyer Assessment of the Upper Extremity (FMAUE) score, we used the definition of Kundert et al. (Kundert et al. 2019):

1. **Initial Impairment (FMAUEii)**: In Eq. 1a, FMAUEii represents the initial impairment, which is calculated by subtracting the initial measurement of the FMAUE (FMAUEi) obtained shortly after the stroke from 66 - the maximum possible score.

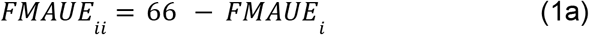
2. **Change in FMAUE (ΔFMAUE)**: Eq. 1b defines ΔFMAUE as the change in the FMA-UE score. It is determined by subtracting the final measurement of the FMA-UE score (FMAUEf) from the initial impairment score (FMAUEii).

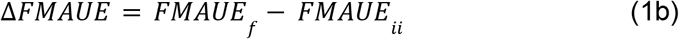
3. **Proportional Recovery Rule (PRR)**: Eq. 1c illustrates the change in FMA-UE according to the proportional recovery rule. This rule posits that at three months post-stroke, patients should recover approximately 70% of their maximum potential improvement, with the slope around 0.7.

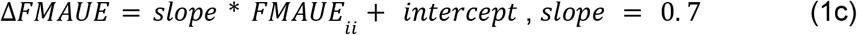

### Introduction of the Mouse Proportional Recovery Rule (MRR)

To assess the applicability of the proportional recovery rule (PRR) in a mouse model, we here introduce the mouse recovery rule (MRR). The PRR, commonly applied in human studies, was evaluated to determine its relevance to behavioral data in mice. To this end, a new rule specific to mice was defined, referred to as the mouse recovery rule (MRR). Several aspects needed to be considered in translating the PRR calculation to the mouse model. First, the behavior tests measure a sensorimotor deficit, thus the logic is reversed compared to the FMA-UE score. Another difference, or even an advantage, is that true baseline behavioral data, i.e., before stroke, are available. Accounting for these considerations, the equations were translated as follows:

1. **Initial Impairment (DSii)**: Eq. 1 uses the difference between the healthy state (FMA-UE ∼66 or ceiling value) and the FMA-UE score shortly after the stroke. This is translated in eq. 2a by replacing the ceiling value of 66 in the FMA-UE score with the baseline value (DS_BL_) of the deficit score, representing the healthy state, and the deficit score on day 3 (DS_P3_) post-stroke, representing the deficit in the (sub-)acute phase.
2. **Change in Deficit Score (ΔDS):** Eq. 2b defines the deficit score as the difference between the initial impairment (DS_ii_) and the deficit score at 28 days post-stroke (DSP_28_), also referred to as the “Change Observed” throughout the text (Fig. 3).
3. **Proportional Recovery Rule (PRR) for the Deficit Score:** In Eq. 2c, the change in the deficit score follows the same logic as Eq. 1c, with a slope and an intercept. Instead of defining a fixed value of 0.7, this value was regressed based on our data.

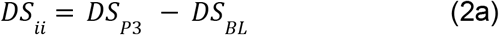

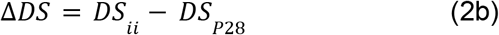

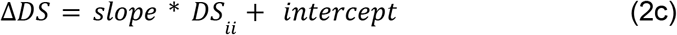

### Iterative cluster analysis

A two-step approach was developed, which consisted of individual regression analysis and iterative clustering refinement. First, the change observed in motor deficit (Eq. 2b) was plotted against the initial impairment (Eq. 2a) for each subject, and a linear fit was applied to the data to determine the slope and intercept of the regression line. This approach allowed for identifying recovery patterns similar to those observed in the human Proportional Recovery Rule (PRR) and Mouse Recovery Rule (MRR). Specifically:

1. **Initial Regression and Clustering:** The MRR was derived with a slope and intercept, while the PRR had a fixed slope of 0.7 with an undefined intercept. The intercept from the MRR fit was used to define the PRR line, and both lines were plotted for comparative analysis. Subjects were then initially clustered based on their adherence to these recovery lines. Clustering was performed by calculating the Euclidean distance between each subject’s data point and the two recovery lines (PRR and MRR). Subjects were assigned to the cluster of the line to which they were closer based on this Euclidean distance.
2. **Iterative Refinement:** In the second step, an iterative process was employed to refine the clustering. This process involved recalculating the slope and intercept of the MRR line using only the data points assigned to the MRR cluster. The updated intercept was then applied to redefine the PRR line. The Euclidean distance between each data point and both lines was recalculated, and the clustering was updated iteratively. This refinement process continued until convergence was achieved, meaning clustering assignments stabilized and did not change significantly between iterations.
3. **Outlier Detection and Removal:** Outliers were identified and removed using the interquartile range (IQR) rule to ensure the accuracy of the clustering. This process was applied initially and after each iteration to prevent extreme values from skewing the results.

Utilizing Euclidean distance, this approach ensured that the clustering was based on a comprehensive measure of proximity to the defined recovery lines, reflecting both horizontal and vertical distances. The iterative process allowed for precise refinement of the recovery line parameters and the accurate classification of subjects into the PRR or MRR clusters.

### Statistics

Statistical tests for analyzing the dynamic changes over time with consideration of multiple groups were conducted using *Python 3*.*10*.*7* and the *statsmodels* and *scipy*.*stats* libraries. A parametric mixed model analysis was employed to test the null hypothesis (i.e. no difference between groups or time points). Post-hoc tests were conducted for multiple comparisons: Tukey to identify differences between each time point within each group, and a Šidák to assess differences between groups at specific time points.

### Data and Code Availability

The project dataset containing all data, code, scripts, and documentation according to the FAIR data management workflow (Kalantari et al. 2023) can be accessed online https://gin.g-node.org/Aswendt_Lab/2024_Kalantari_PRR.

## Results

### Variability in Recovery After Stroke: Overall Patterns

The three behavioral tests, rotating beam, cylinder, and grid walk test, showed spontaneous recovery in stroke mice (**Fig. 1**) with considerable individual variability (**Supplementary Fig. S2**). The longitudinal data was assessed over time and between groups using a mixed-effects model approach (**Supplementary Table S1**). The tests differed in their sensitivity to detect differences between stroke and sham groups at selected time points. The rotating beam test primarily distinguished stroke from sham mice in the acute phase (day 3 p<0.001, day 7 p<0.01), while the cylinder test, grid walk test, and deficit scores all showed differences between stroke and sham animals up to the chronic phase (days 3, 7, 21, and 28 p<0.001). Regarding time differences, the deficit scores, rotating beam test, cylinder test, and grid walk test revealed significant deficits up to day 28 post-stroke compared to baseline in the stroke group (p<0.001), with the exception of the rotating beam test (only up to day 21 with p<0.001). For the sham group, no significant differences were observed in any of the tests or the deficit scores between any of the time points and baseline.

**Figure 1:**
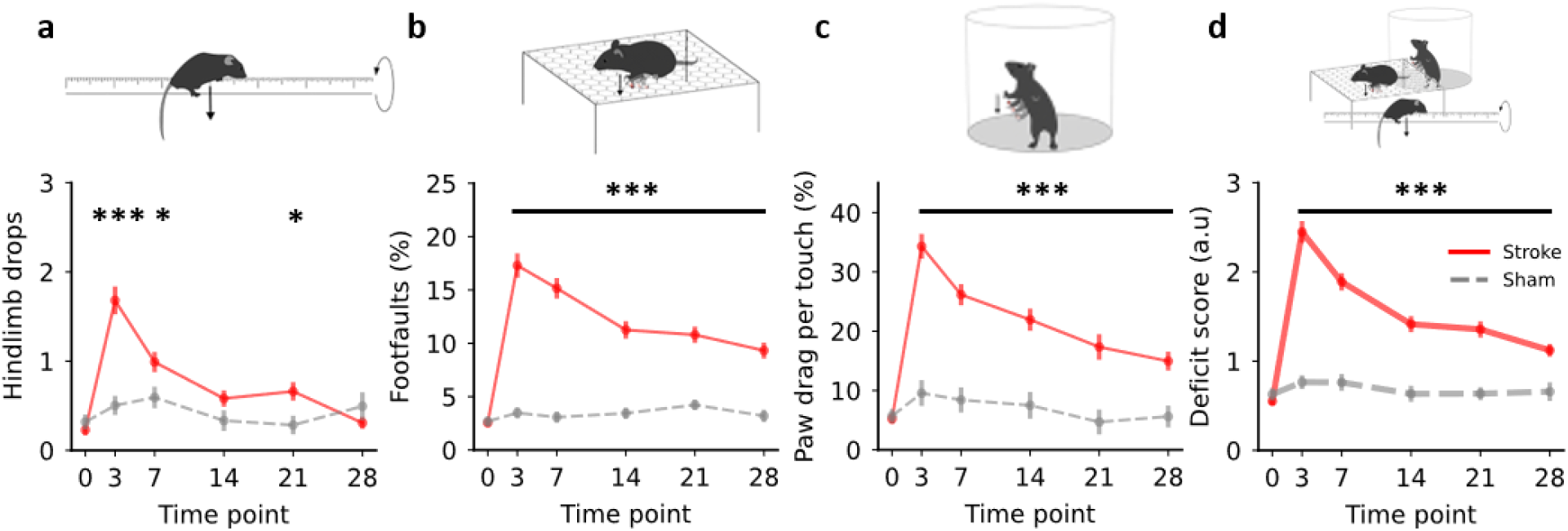
Dynamics of spontaneous amelioration of specific motor deficits four weeks after cortical stroke. (a) Rotating beam tests: number of hindlimb drops recorded when the mouse walks over a rotating beam towards the homecage. (b) Grid walk test: number of footfaults related to the total number of steps during spontaneous walking on an elevated grid. (c) Cylinder test: number of paw drags per touch with the affected paw at the cylinder wall. (d) Deficit score, calculated by averaging the z-transformed values of each test (a-c). Data in graphs are shown as mean ± SEM. Significant differences between stroke and sham at the same time points are shown as *p ≤ 0.05, **p ≤ 0.01, and ***p ≤ 0.001, respectively. Mixed model analysis was used to assess significance, followed by post hoc Tukey tests for within-group time point comparisons and Šidák multiple comparison tests for between-group significant time point differences.

### Iterative cluster analysis

To analyze the PRR, the z-transformed values of the three tests were used as the deficit score to plot the change observed between 28 days post-stroke (P28) and initial impairment (Eq. 2b) versus the initial impairment itself (Eq. 2a). The linear fit changed with each iteration until convergence between the 4th and 5th iteration. Further interactions did not change the slope or the intercept for the MRR fit further (**Fig. 2**). The intercept for both clusters converged at -0.72. The goodness of fit for both MRR and PRR was high (R-squared of 0.93 and 0.80, respectively). To determine if the slopes between the PRR and MRR were significantly different, an F-test was performed. The results showed that the overall slopes were not identical, with an F-value of 4.2 (DFn = 1, DFd = 91) and p<0.05 (**Supplementary Table 3**).

**Figure 2:**
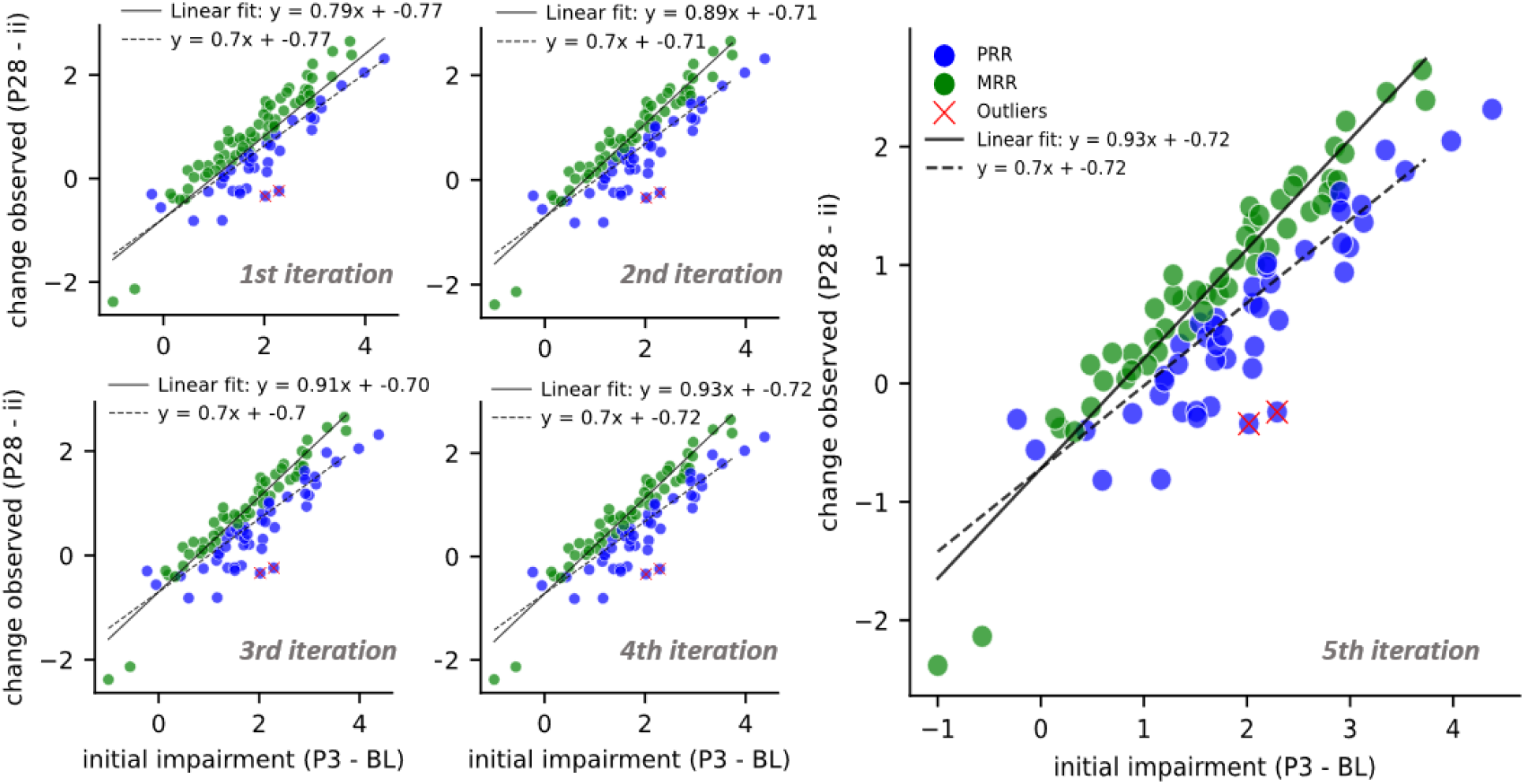
Iterative cluster analysis process for the mouse recovery rule (MRR) and proportional recovery rule (PRR). For the calculated deficit scores, the change observed (difference between 28 days poststroke and initial impairment) for each subject was plotted against their initial impairment (difference between Acute phase or P3 and baseline values), and a linear fit was applied to determine the slope and intercept for the MRR. The PRR was assigned a fixed slope of 0.7, with the intercept derived from the MRR fit. Subjects were initially categorized into two clusters based on their proximity to the PRR or MRR line. An iterative process refined clustering by recalculating linear fits and updating cluster assignments until convergence. The figure illustrates the convergence of this iterative process. By the 5th iteration, the slope and intercept of the linear fit stabilized at 0.93 and -0.72, respectively. This stability indicates no further changes in these parameters, confirming the consistency of cluster assignments. The goodness of fit for the MRR and PRR was evaluated, resulting in an R-squared of 0.9316 for the MRR fit and 0.8071 for the PRR fit. This analysis was performed on a cohort of 38 subjects, 22 of whom were assigned to the MRR group and 16 to the PRR group.

The initial cohort consisted of 125 subjects, however, the final analysis was performed on 95 mice due to the exclusion of subjects where critical time points were missing. Specifically, missing data on time points needed for the calculation of the initial impairment and the observed change led to this reduction in animal numbers. The final clustering resulted in 47.36% (N=45) mice in the PRR and 52.64% (N=50) in the MRR cluster. Mice in the PRR cluster exhibited the predefined fixed slope of 0.7, while mice in the MRR cluster displayed a slope of 0.93, indicative of a distinct recovery trajectory.

### Variability in Recovery After Stroke: Cluster-Specific Trajectories

Clustered behavior data were statistically compared over time and between groups using a mixed-effects model approach (**Supplementary Table S2**). Both the PRR and MRR clusters showed reduced deficits across all behavioral tests, with significant improvements noted from day 0 to day 3, day 7, day 14, day 21, and day 28 post-stroke across all tests (p<0.01), except for the rotating beam test in the PRR cluster from day 0 to day 28, which showed no significant recovery (**Fig. 3)**. The difference in recovery between day 21 and day 28 was less pronounced, showing a significant change only in the MRR cluster across cylinder tests and deficit scores (p<0.05), suggesting variable recovery trajectories between clusters (**Supplementary Fig. 2)**. PRR and MRR started from similar deficit levels (no significant differences at day 3), however, MRR mice recovered faster and more thorougly, as the comparison with PRR-fitting mice was significant at 14 (p<0.05) and up to 29 days (p<0.001). Except for the grid walk test (p<0.001), the MRR deficit was not significantly different to sham mice at 28 days, In contrast, the PRR group remained with a significantly higher deficit compared to sham mice (p<0.001, except for the rotating beam test with p>0.05).

**Figure 3:**
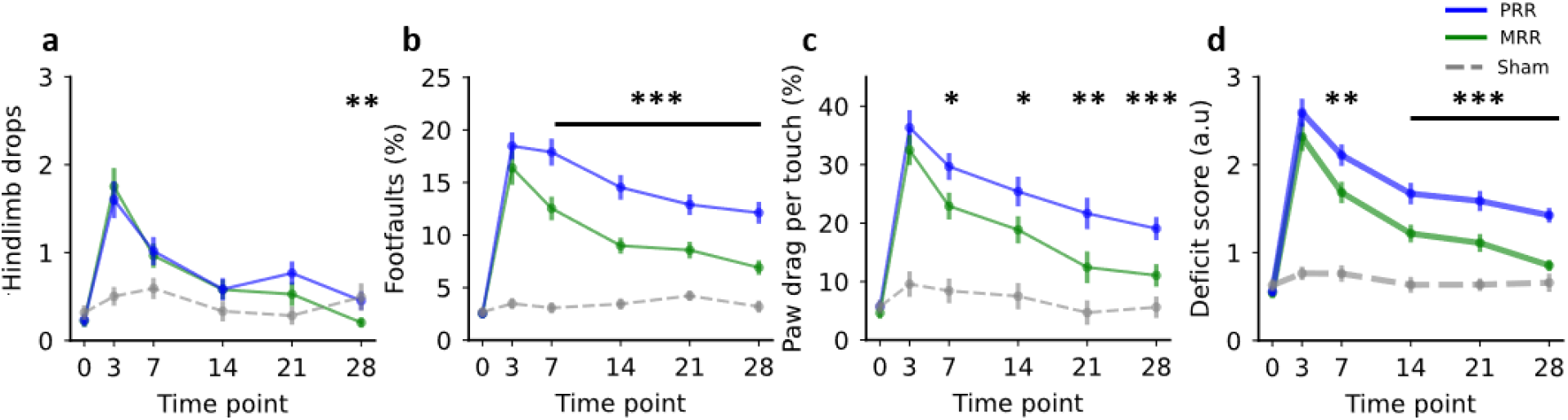
Dynamics of behavioral test performance over time for each group after cluster assignments. (a) Rotating beam test: number of hind limb drops recorded for a walking task on a rotating beam. (b) Grid walk test: number of footfaults calculated as a percentage of total footsteps. (c) Cylinder test: number of paw drags, calculated as a percentage per touch with the affected paw at the cylinder wall. (d) Deficit score, calculated by averaging the z-transformed values of each test. Data are shown as mean ± SEM. Significant differences between the two cluster groups (PRR and MRR) at the same time points are indicated by *p ≤ 0.05, **p ≤ 0.01, and ***p ≤ 0.001. Mixed model analysis was used to assess significance, followed by post hoc Tukey tests for within-group time point comparisons and Šidák multiple comparison tests for between-group significant time point differences.

### Role of stroke lesion size and location

To verify that the differences in functional recovery were not driven by lesion size or location, a comprehensive qualitative voxel- and quantitative atlas-based analysis of the stroke lesion as determined using T2-weighted MRI was conducted (**Fig. 4**). The semi-automatically defined lesion masks were registered to the Allen Mouse Brain Atlas (**Fig. 4a**). The qualitative voxel-wise comparison showed no differences between the MRR and PRR cluster compared to the combined group in terms of lesion pattern and lesion extent (**Fig. 4b**). This was also reflected in a similar lesion volume for the MRR (15.32+-10.61) and PRR (18.57+-10.47) group (**Fig. 4c**). Further, there was no difference in the lesion location as quantified by the percentage of lesion mask overlap with brain atlas regions (**Fig. 4d)**.

**Figure 4.**
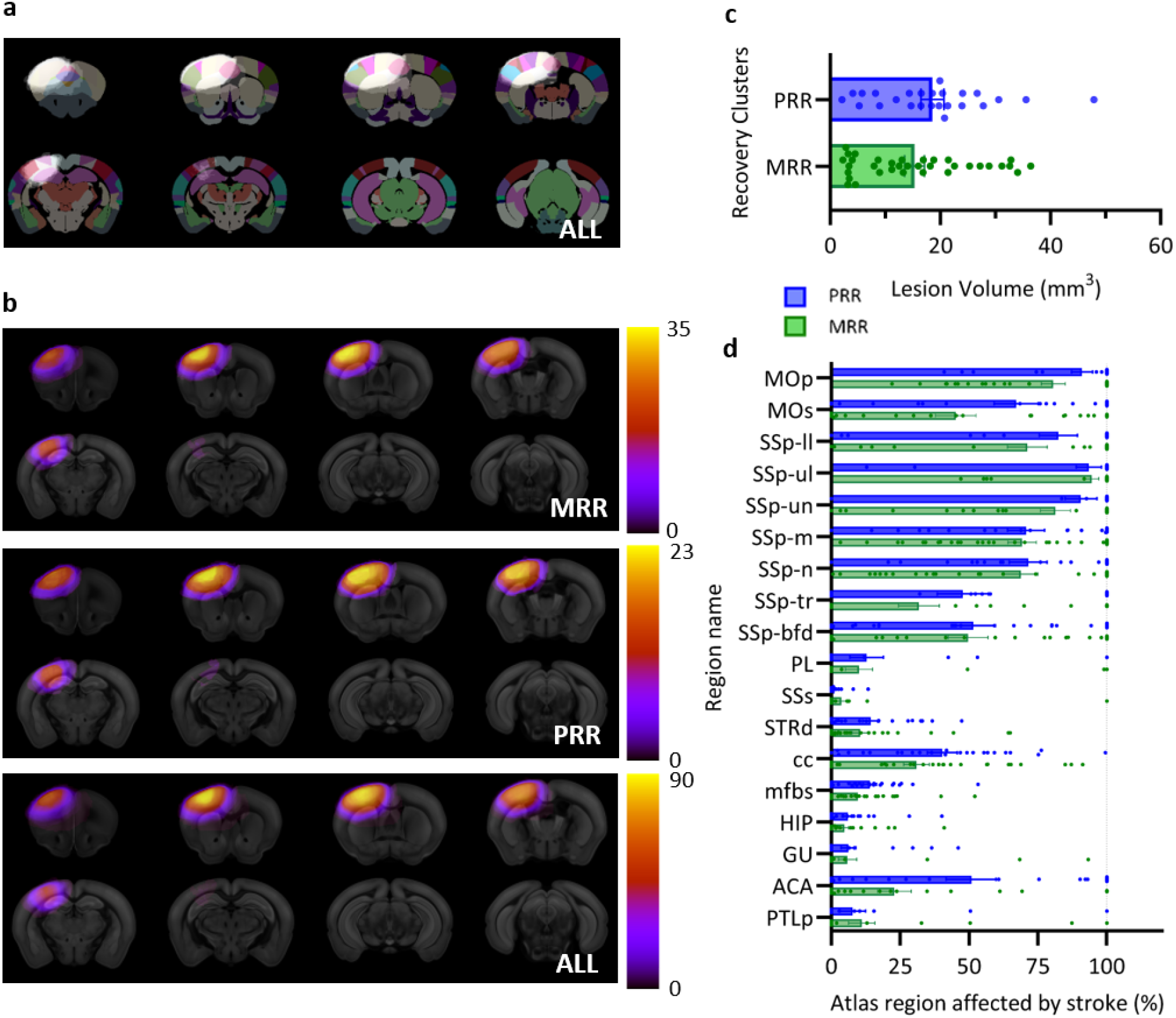
The role of stroke lesion size, and location. (a) Stroke incidence map and modified Allen Brain Atlas (48 regions) overlay. (b) Incidence maps of cortical strokes for MRR, PRR and all mice showing no visual difference. (c) No lesion volume differences between the two recovery clusters MRR and PRR (unpaired t-test, p > 0.05). (d) There was no difference in the stroke infarct area per atlas region between the two recovery clusters MRR and PRR (mixed model analysis with subsequent Sidak multiple comparisons, p > 0.05).

The clustering based on the linear fit approach (**Fig. 2d**) contained two outliers as detected based on the interquartile range rule. These outliers showed a relatively high initial impairment and a negative change observed, i.e., poor improvement over time. MRI data was available for only one of them, which revealed a stroke volume of approximately 28.36 mm^3^. This is notably larger than the average stroke volume of 16.31±10.69 mm^3^ observed across all data points in both clusters. Additionally, the two lowest points in the MRR cluster, in terms of the *change observed* value, could be considered outliers. However, based on the interquartile range rule, this was not the case. These subjects exhibited lesion sizes of 32.6 mm^3^ and 36.4 mm^3^, respectively, again indicating relatively large strokes.

## Discussion

We present a new approach combining three different motor tests to improve the reliability of motor assessment in mice and to compensate for differences in sensitivity over a longer time frame (Balkaya and Cho 2019). The results revealed that approximately 40% of the mice with cortical strokes followed the PRR and 60% followed a new recovery trajectory with a slope of approximately 0.9, which we termed the mouse recovery rule (MRR). MRR-fitting mice recover 90% of their initial impairment within the first four weeks post-stroke. Notably, the different recovery trajectories were not related to lesion size or location differences. This study provides new insights into the translational applicability of the PRR to mouse studies and emphasizes the individual differences in recovery mechanisms present beyond lesion size and location.

### Classification of recovery trajectories

Identifying recovery patterns after stroke is crucial for advancing rehabilitation strategies, with the Proportional Recovery Rule (PRR) representing one established approach (Prabhakaran et al. 2008). Expanding on the classification of recovery trajectories, Van der Vliet et al developed a longitudinal mixture model using an exponential recovery function (van der Vliet et al. 2020), challenging the PRR’s assumptions and addressing criticisms of its predictive oversimplification. Furthermore, Nashet’s application of a K-means clustering approach on non-human primate stroke data (Nashed et al. 2024) might provide a deeper separation of the various sources of different recovery patterns. Additionally, Jeffers et al. adapted the PRR to rodent models (Jeffers et al. 2018), categorizing subjects into ‘fitters,’ who follow the PRR, ‘non-fitters,’ who do not adhere to the PRR, and ‘decliners,’ who do not recover at all. This categorization demonstrates the PRR’s applicability across species and underscores the variability in recovery trajectories. In our study, we observed similar classifications, with “non-fitters” termed as MRR exhibiting better outcomes than the PRR group, which aligns with the previously defined ‘fitters’. In contrast to Jeffers’ study where ‘non-fitters’ experienced poorer recovery, the MRR group showed significantly better recovery outcomes. This observation led to the proposal of the MRR as a novel predictive model tailored to mouse recovery post-stroke.

### Behavior tests to assess spontaneous recovery after stroke

In line with previous studies, mice with cortical strokes induced by photothrombosis recovered substantially during the first weeks after stroke (Clarkson et al. 2013; Götz et al. 2023; Yu et al. 2015). As expected, there were considerable differences in the recovery rate and the sensitivity in detecting long-term motor deficits, which can be attributed to the different weighting ranging from general motor coordination (e.g., gait) to detailed motor output (e.g., correct paw placement on the grid). The results revealed that the rotating beam test is highly susceptible to early balance and coordination deficits but less so for long-term ones. In contrast, the cylinder test demonstrated a more gradual recovery, with significant differences between the stroke and sham groups observed at multiple time points, indicating its sensitivity to persistent forelimb use asymmetry. The grid walk test, assessing foot faults, revealed the highest sensitivity to sensorimotor impairments, with significant differences between the stroke and sham groups maintained until 3 weeks post-stroke, highlighting its efficacy in detecting acute and chronic deficits in motor coordination and proprioception.

The combination of these behavioral tests, which all showed a significant impairment at 4 weeks compared to baseline, allowed us to rule out a primary concern that most of the behavioral tests lose sensitivity within weeks so that the PRR cannot hold for animal models of stroke (Balkaya and Cho 2019). In contrast, our clustering analysis revealed that a substantial number of mice (40%) adhere to the proportional recovery rule, suggesting that this rule may have relevance beyond clinical settings. Compared to the only previous rat MCAO study (Jeffers et al. 2018), which found that 30% of the rats fit the PRR rule, our findings underscore that a similar proportion of mice in our study adhere to the PRR. The identification of a distinct cluster with its own regression pattern highlights the heterogeneity of post-stroke recovery and underscores the importance of individualized treatment approaches.

### Role of stroke lesion size and location

Experimental studies of stroke recovery have extensively explored the role of lesion size, location, and neural circuit integrity. Lesion size is a common imaging marker to predict or explain stroke outcomes (Sperber et al. 2023). However, as shown in human and rodent studies, the additional analysis of the lesion topology improves the prognostic value (Knab et al. 2023; Scheulin et al. 2021; Tscherpel et al. 2024). Many rodent models use large infarcts, which may not accurately reflect the typical, smaller strokes experienced by humans. This discrepancy can impact the relevance of findings to human stroke conditions. In large population studies the lesion size ranges from 28 to 80 cm^3^, which translates to approximately 4.5% to 14% of the total volume of the ipsilesional hemisphere (Carmichael 2005). We used photothrombosis to induce homogeneous and well-defined lesions. Our data show a mean of 7±2.21% for the PRR cluster and 5.6±2.02% for the MRR cluster. This aligns with the retrospective analysis of studies using endothelin-1 to induce forelimb sensorimotor cortex and dorsolateral striatum lesions in rats (Jeffers et al. 2018). Approximately 30% of the rats were identified as fitters of the PRR, which had smaller lesions (5.3%) than the rats that did not show proportional recovery or deteriorated over time (8.3%). Given a comparable lesion volume, the ratio of PRR fitters differs between up to 80% (humans), 30% (rats), and 47% (mice). Although the lesion volume overlaps across species, the lesion location differs and thus the existing data cannot be directly compared. Effective recovery modeling must consider specific neuronal circuits affected by the stroke (Carmichael 2005). Byblow et al. studied the role of the integrity of the corticomotor pathway in proportional recovery. They discovered that patients with a viable ipsilesional corticomotor pathway were more likely to experience proportional recovery, while those with compromised pathways exhibited reduced recovery (Byblow et al. 2015). Such a longitudinal characterization of motor fiber integrity using diffusion MRI in rodents is missing. Future studies may include a comprehensive analysis of lesion topology and the differential effects on functional circuits and structural brain connectivity.

### Transformation of behavioral to deficit scores

The process of deriving a deficit score through z-transformation involves converting individual scores into z-scores, and integrating these scores to provide a comprehensive measure of motor recovery. This standardized deficit score was analyzed longitudinally and across experimental groups (sham and stroke) to evaluate the impact of stroke and subsequent recovery. This method facilitates a standardized comparison of motor function across different tests and time points, which is crucial for capturing the behavioral recovery trajectory in stroke-affected mice. Using z-transformation is particularly pertinent because this statistical transformation is typically suitable for data with a normal distribution. However, in this study, a normal distribution across the entire cohort of stroke mice was not assumed. Initially, stroke mice scores were expected to be comparable to sham mice at baseline, but immediately post-stroke, a significant deficit was anticipated, leading to a deviation from normal distribution. While normal distribution within each time point is ideal, factors such as variability in test sensitivity and the subjective nature of assessing the quantified values of each test may challenge this assumption. Despite these complexities, merging the results from three tests into a single deficit score helps mitigate the limitations of individual tests by leveraging their combined strengths (Guilloux et al. 2011). This approach is also supported by the finding that the deficit score revealed more robust significant differences. In light of dynamic changes observed from baseline to the chronic phase, applying the z-transformation across the entire cohort is crucial. This approach allows for meaningful comparisons among subjects with varying scores at different tests and ensures consistency across different time points. Applying z-transformation separately for each time point would obscure these longitudinal dynamics, underscoring the importance of a comprehensive transformation approach. Given the variability in behavioral data and the need to avoid fixed boundaries (Bowman et al. 2021) that might constrain interpretation, the z-score transformation was ultimately chosen for its ability to standardize the data while maintaining flexibility. This decision and its implications are further discussed in the subsequent section.

### Statistical considerations

#### Coupling effects and measurement scales

Mathematical coupling can be a significant issue in PRR-related statistical analyses (Bowman et al. 2021; Chong, Wang, and Stinear 2023). Others argue that when properly analyzed, the coupling can be viewed as a notational construct rather than a true confound (Chong et al. 2023). Likewise, our analysis might be subjective to mathematical coupling due to the definitions of the initial impairment (deficit at day 3 - deficit at baseline) and the change in deficit score (deficit at day 28 - initial impairment). Future studies may mitigate the effects of mathematical coupling by using alternative modeling approaches that do not rely on the change variable (Y - X) (Bowman et al. 2021), and fitting models to raw performance scores over time (van der Vliet et al. 2020).

Another confound in PRR-related statistics is the ceiling effect of the maximum FM-UE score of 66, which can compress scores at the upper end and skew the results (Bowman et al. 2021). Discarding or transferring data at the ceiling was suggested to reduce bias. In our case, the deficit score is an average of three behavioral assessments, each of which underwent z-transformation to enable averaging. As the z-transformation creates no ceiling or floor effect, this approach eliminates the problem of scaling thresholds. This perspective is crucial for our study, as it suggests that the theoretical concerns raised by mathematical coupling and ceiling effects do not fundamentally undermine the validity of our observed correlations. By carefully addressing these concerns using z-scores and considering the limitations of scales, we ensure the robustness of our analysis and the validity of applying the proportional recovery rule in our mouse model of stroke recovery.

#### Availability of baseline data

In human studies, the Fugl-Meyer Assessment of the Upper Extremity (FMA-UE) score reaches a maximum possible score of 66 serving as a reference point for initial impairment calculations. However, in our mouse model, we leverage baseline behavioral data obtained before stroke inducement. This allowed a more precise measurement of the initial impairment (DSii), as the actual pre-stroke performance of each mouse could be used rather than an assumed ceiling value, which can help mitigate the coupling effect as it increases variability in the initial impairment making it dependent on two variables.

### Limitations and Future Directions

While our clustering approach provides valuable insights, it is essential to acknowledge its limitations, including the missing verification in an aged and sex-matched cohort of mice, and the potential for variability across stroke models. It has been stressed that the preclinical stroke research should include biological variables and clinically relevant features, such as age, sex, diabetes, obesity, and hypertonia (Corbett et al. 2017; Wolf and Ergul 2021). One limitation of our behavior tests is the measure of single partially overlapping deficits, which are more prone to compensation rather than a comprehensive score reflecting true recovery (Prabhakaran et al. 2008). However, all tests reflect spontaneous behavior, which is in general less prone to compensatory effects (Corbett et al. 2017). To our knowledge, no comprehensive scoring system has complemented the FMA-UE scoring system developed for mice. The Bederson scale, helpful for distinguishing stroke lesion size in the acute phase, has not been validated to predict recovery outcomes (Bieber et al. 2019). Recently, deep behavioral phenotyping with kinematic analysis has shown movement patterns related to compensation, like adapting contralateral limb stability and reducing trunk distance from the target prior to reaching, which are common in human stroke patients (Balbinot et al. 2018; Weber et al. 2022). Future studies would benefit from a more detailed translational kinematic assessment. Also increasing the sample size would allow alternative clustering methods, such as the longitudinal mixture model of FMA-UE recovery (van der Vliet et al. 2020).

### Conclusion

In summary, our study demonstrates the translation of the proportional recovery rule from clinical practice to a mouse stroke model. By employing a clustering approach, we delineate distinct recovery trajectories, that are not driven by lesion size or location. These findings suggest that alternative recovery mechanisms drove post-stroke recovery in this homogeneous group of mice. The findings contribute to our understanding of stroke pathophysiology and may inform the development of threrapeutic interventions later to be translated to stroke patients.

## Acknowledgments

We acknowledge Frederique Wieters, Veronika J. Fritz, Olivia Käsgen, Niklas Palast, Marc Schneider, and Victor Vera Frazao for their contributions in acquiring and processing data for this study.

## Sources of Funding

This work was funded by the Friebe Foundation (T0498/28960/16) and the Deutsche Forschungsgemeinschaft (DFG, German Research Foundation) – Project-ID 431549029 – SFB 1451.

## Disclosures

The authors declare that they have no conflict of interest.

## Notes

https://gin.g-node.org/Aswendt_Lab/2024_Kalantari_PRR

### Competing Interest Statement

The authors have declared no competing interest.

