## Supplement for "Proportional recovery in mice with cortical stroke"

### Supplementary Material

All coding, processing, and further analysis started with 125 stroke and 35 sham mice. Among the 125 stroke animals, 67 animals had behavioral data without accompanying MRI, while 58 animals had both behavioral and MRI data. MRI was performed and used to analyze lesion size and location in 90 stroke mice; however, 32 had either incomplete behavioral data or no behavioral data. In total, 58 stroke animals had both behavior and MRI information for comprehensive analysis (**Supplementary Fig. S1**).

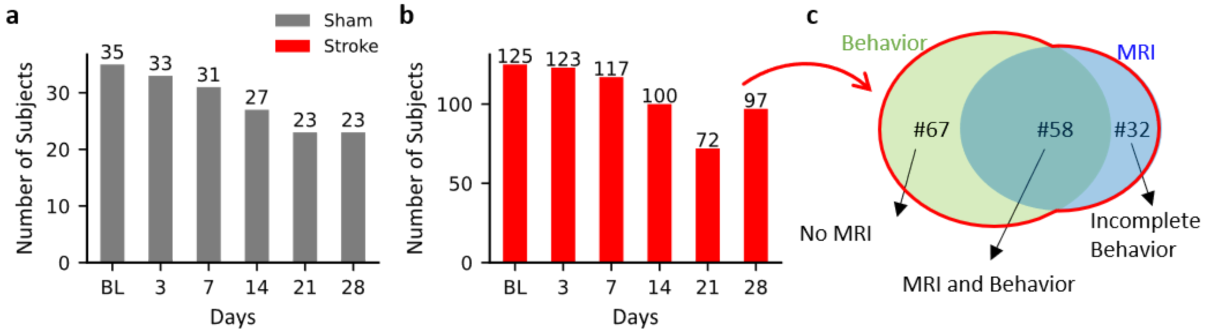

**Figure S1: Overview of data collection, and availability for stroke and sham mice.** Details of data collection for (a) n=35 sham mice and (b) n=125 stroke mice. Behavioral assessments were performed at baseline and post-stroke on days 3, 7, 14, 21, and 28. Lesion size and location were determined using in vivo MRI on days 3 or 7 post-surgery. (c) Distribution of available data for stroke mice of the 125 mice, with 58 having both behavioral and MRI data, and 67 having only behavioral data, taking the initial 125 of the BL as reference. MRI data were obtained for 90 mice, but 32 had incomplete or no behavioral data, as indicated.

Photothrombotic stroke in the sensorimotor cortex was induced in the n=125 mice and sham surgery was applied in the n=35 mice as described before (Götz et al. 2023). Briefly, mice were anesthetized with 4% isoflurane and secured in a stereotactic frame with continuous 2% isoflurane. Temperature was monitored and controlled using a rectal probe and heating pad. After shaving and disinfecting the surgical area, a 1.5 cm incision was made, and the exposed skull was cleaned. A 561 nm laser was aligned to the primary motor cortex (M/L: 2.00 mm and A/P: 0.00 mm), and mice received an intraperitoneal injection of 1000 µg Rose Bengal dye. After 5 minutes, the laser was applied through the skull for 15 minutes at 50 mW. Sham surgery followed the same steps but without laser exposure. The wound was closed post-surgery, and mice recovered in a heated chamber. Analgesia included Tramadol (1 mg/ml) in the drinking water for 3 days before and after surgery, and an intraoperative injection of 4 mg/kg Caprofen. Lesion location and size were determined by an in vivo T2-weighted TurboRare sequence as described before (Aswendt et al. 2021) on day 3 or 7 post-surgery and analyzed using the in-house software AIDAmri (Pallast et al. 2019). Motor recovery was measured at day 0 (before stroke = baseline), and 3, 7, 14, 21, and 28 days post-stroke using the rotating beam, cylinder, and grid walk test (Aswendt et al. 2022). The experimenters were blinded to the animal group (stroke/sham) and the experimenter who performed the stroke surgeries did not perform the

MRI and behavior tests. Unbiased and blinded documentation was performed in an electronic database (Pallast et al. 2018).

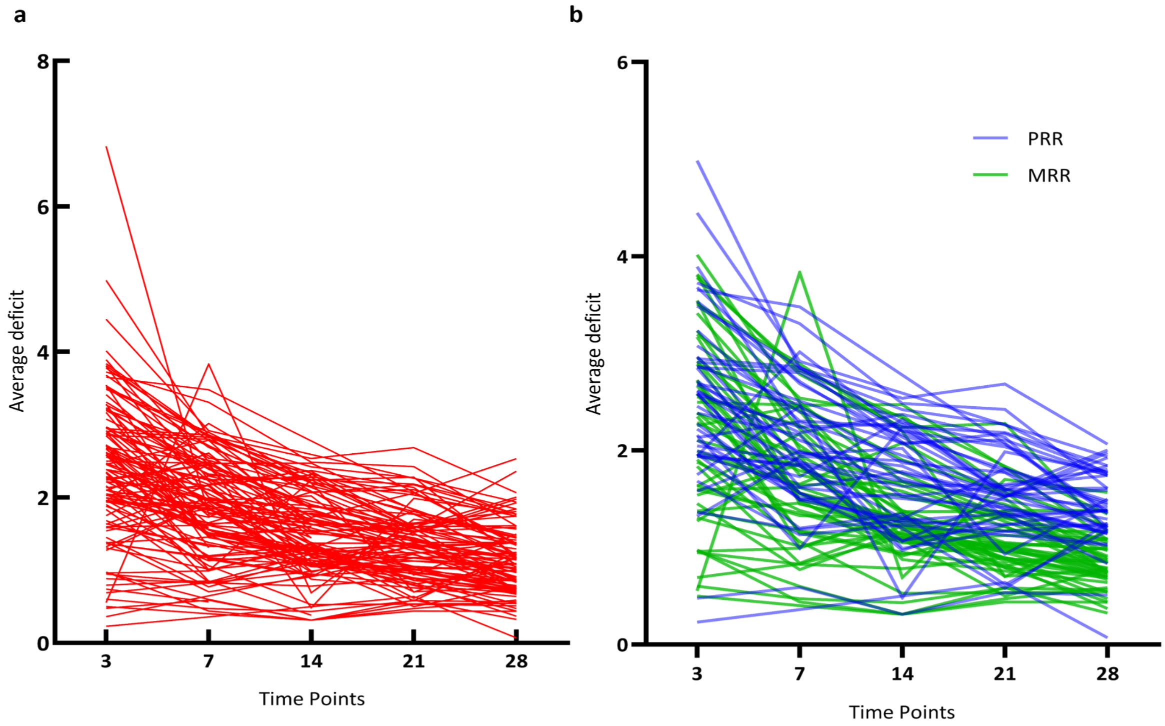

**Figure S2: Individual reduction of motor deficit before and after clustering.** Spaghetti plots for the average deficit of (a) all stroke animals (b) PRR (blue) and MRR (green) mice between 3 and 28 days post-stroke.

Table S1: Detailed statistics of Figure 2 in the main text

| Paw drags | | | Hindlimb drop | | | Foot faults | | | Deficit score | | |
|---|---|---|---|---|---|---|---|---|---|---|---|
| *Mixed-effect analysis* | | | *Mixed-effect analysis* | | | *Mixed-effect analysis* | | | *Mixed-effect analysis* | | |
| **Comparison** | **Group** | **Sig.** | **Comparison** | **Group** | **Sig.** | **Comparison** | **Group** | **Sig.** | **Comparison** | **Group** | **Sig.** |
| Time | all | *** | Time | all | *** | Time | all | *** | Time | all | *** |
| Group | all | *** | Group | all | *** | Group | all | *** | Group | all | *** |
| Interaction | all | *** | Interaction | all | *** | Interaction | all | *** | Interaction | all | *** |
| *Multiple comparison test: Tukey* | | | *Multiple comparison test: Tukey* | | | *Multiple comparison test: Tukey* | | | *Multiple comparison test: Tukey* | | |
| **Comparison** | **Group** | **Sig.** | **Comparison** | **Group** | **Sig.** | **Comparison** | **Group** | **Sig.** | **Comparison** | **Group** | **Sig.** |
| 0 vs.3 | Stroke | *** | 0 vs.3 | Stroke | *** | 0 vs.3 | Stroke | *** | 0 vs.3 | Stroke | *** |
| 0 vs.7 | Stroke | *** | 0 vs.7 | Stroke | *** | 0 vs.7 | Stroke | *** | 0 vs.7 | Stroke | *** |
| 0 vs.14 | Stroke | *** | 0 vs.14 | Stroke | *** | 0 vs.14 | Stroke | *** | 0 vs.14 | Stroke | *** |
| 0 vs.21 | Stroke | *** | 0 vs.21 | Stroke | *** | 0 vs.21 | Stroke | *** | 0 vs.21 | Stroke | *** |
| 0 vs.28 | Stroke | *** | 0 vs.28 | Stroke | ns | 0 vs.28 | Stroke | *** | 0 vs.28 | Stroke | *** |
| 0 vs.3 | Sham | ns | 0 vs.3 | Sham | ns | 0 vs.3 | Sham | ns | 0 vs.3 | Sham | ns |
| 0 vs.7 | Sham | ns | 0 vs.7 | Sham | ns | 0 vs.7 | Sham | ns | 0 vs.7 | Sham | ns |
| 0 vs.14 | Sham | ns | 0 vs.14 | Sham | ns | 0 vs.14 | Sham | ns | 0 vs.14 | Sham | ns |
| 0 vs.21 | Sham | ns | 0 vs.21 | Sham | ns | 0 vs.21 | Sham | ns | 0 vs.21 | Sham | ns |
| 0 vs.28 | Sham | ns | 0 vs.28 | Sham | ns | 0 vs.28 | Sham | ns | 0 vs.28 | Sham | ns |
| *Multiple comparison test: Šidák* | | | *Multiple comparison test: Šidák* | | | *Multiple comparison test: Šidák* | | | *Multiple comparison test: Šidák* | | |
| **Comparison** | **Group** | **Sig.** | **Comparison** | **Group** | **Sig.** | **Comparison** | **Group** | **Sig.** | **Comparison** | **Group** | **Sig.** |
| Stroke vs.Sham | TP 0 | ns | Stroke vs.Sham | TP 0 | ns | Stroke vs.Sham | TP 0 | ns | Stroke vs.Sham | TP 0 | ns |
| Stroke vs.Sham | TP 3 | *** | Stroke vs.Sham | TP 3 | *** | Stroke vs.Sham | TP 3 | *** | Stroke vs.Sham | TP 3 | *** |
| Stroke vs.Sham | TP 7 | *** | Stroke vs.Sham | TP 7 | ** | Stroke vs.Sham | TP 7 | *** | Stroke vs.Sham | TP 7 | *** |
| Stroke vs.Sham | TP 14 | *** | Stroke vs.Sham | TP 14 | ns | Stroke vs.Sham | TP 14 | *** | Stroke vs.Sham | TP 14 | *** |
| Stroke vs.Sham | TP 21 | *** | Stroke vs.Sham | TP 21 | ** | Stroke vs.Sham | TP 21 | *** | Stroke vs.Sham | TP 21 | *** |
| Stroke vs.Sham | TP 28 | *** | Stroke vs.Sham | TP 28 | ns | Stroke vs.Sham | TP 28 | *** | Stroke vs.Sham | TP 28 | *** |

#### Table S2: Detailed statistics of Figure 4 in the main text

| Paw drags | | | Hindlimb drop | | | Foot faults | | | Deficit score | | |
|---|---|---|---|---|---|---|---|---|---|---|---|
| *Mixed-effect analysis* | | | *Mixed-effect analysis* | | | *Mixed-effect analysis* | | | *Mixed-effect analysis* | | |
| **Comparison** | **Group** | **Sig.** | **Comparison** | **Group** | **Sig.** | **Comparison** | **Group** | **Sig.** | **Comparison** | **Group** | **Sig.** |
| Time | all | *** | Time | all | *** | Time | all | *** | Time | all | *** |
| Group | all | *** | Group | all | ** | Group | all | *** | Group | all | *** |
| Interaction | all | *** | Interaction | all | *** | Interaction | all | *** | Interaction | all | *** |
| *Multiple comparison test: Tukey* | | | *Multiple comparison test: Tukey* | | | *Multiple comparison test: Tukey* | | | *Multiple comparison test: Tukey* | | |
| **Comparison** | **Group** | **Sig.** | **Comparison** | **Group** | **Sig.** | **Comparison** | **Group** | **Sig.** | **Comparison** | **Group** | **Sig.** |
| 0 vs.3 | PRR | *** | 0 vs.3 | PRR | *** | 0 vs.3 | PRR | *** | 0 vs.3 | PRR | *** |
| 0 vs.7 | PRR | *** | 0 vs.7 | PRR | *** | 0 vs.7 | PRR | *** | 0 vs.7 | PRR | *** |
| 0 vs.14 | PRR | *** | 0 vs.14 | PRR | ** | 0 vs.14 | PRR | *** | 0 vs.14 | PRR | *** |
| 0 vs.21 | PRR | *** | 0 vs.21 | PRR | ** | 0 vs.21 | PRR | *** | 0 vs.21 | PRR | *** |
| 0 vs.28 | PRR | *** | 0 vs.28 | PRR | ns | 0 vs.28 | PRR | *** | 0 vs.28 | PRR | *** |
| 0 vs.3 | MRR | *** | 0 vs.3 | MRR | *** | 0 vs.3 | MRR | *** | 0 vs.3 | MRR | *** |
| 0 vs.7 | MRR | *** | 0 vs.7 | MRR | *** | 0 vs.7 | MRR | *** | 0 vs.7 | MRR | *** |
| 0 vs.14 | MRR | *** | 0 vs.14 | MRR | ** | 0 vs.14 | MRR | *** | 0 vs.14 | MRR | *** |
| 0 vs.21 | MRR | * | 0 vs.21 | MRR | * | 0 vs.21 | MRR | *** | 0 vs.21 | MRR | *** |
| 0 vs.28 | MRR | ** | 0 vs.28 | MRR | ns | 0 vs.28 | MRR | *** | 0 vs.28 | MRR | *** |
| *Multiple comparison test: Šídák* | | | *Multiple comparison test: Šídák* | | | *Multiple comparison test: Šídák* | | | *Multiple comparison test: Šídák* | | |
| **Comparison** | **Group** | **Sig.** | **Comparison** | **Group** | **Sig.** | **Comparison** | **Group** | **Sig.** | **Comparison** | **Group** | **Sig.** |
| PRR vs. MRR | TP 0 | ns | PRR vs. MRR | TP 0 | ns | PRR vs. MRR | TP 0 | ns | PRR vs. MRR | TP 0 | ns |
| PRR vs. Sham | TP 0 | ns | PRR vs. Sham | TP 0 | ns | PRR vs. Sham | TP 0 | ns | PRR vs. Sham | TP 0 | ns |
| MRR vs. Sham | TP 0 | ns | MRR vs. Sham | TP 0 | ns | MRR vs. Sham | TP 0 | ns | MRR vs. Sham | TP 0 | ns |
| PRR vs. MRR | TP 3 | ns | PRR vs. MRR | TP 3 | ns | PRR vs. MRR | TP 3 | ns | PRR vs. MRR | TP 3 | ns |
| PRR vs. Sham | TP 3 | *** | PRR vs. Sham | TP 3 | *** | PRR vs. Sham | TP 3 | *** | PRR vs. Sham | TP 3 | *** |
| MRR vs. Sham | TP 3 | *** | MRR vs. Sham | TP 3 | *** | MRR vs. Sham | TP 3 | *** | MRR vs. Sham | TP 3 | *** |
| PRR vs. MRR | TP 7 | ns | PRR vs. MRR | TP 7 | ns | PRR vs. MRR | TP 7 | ** | PRR vs. MRR | TP 7 | ns |
| PRR vs. Sham | TP 7 | *** | PRR vs. Sham | TP 7 | ns | PRR vs. Sham | TP 7 | *** | PRR vs. Sham | TP 7 | *** |
| MRR vs. Sham | TP 7 | *** | MRR vs. Sham | TP 7 | ns | MRR vs. Sham | TP 7 | *** | MRR vs. Sham | TP 7 | *** |
| PRR vs. MRR | TP 14 | ns | PRR vs. MRR | TP 14 | ns | PRR vs. MRR | TP 14 | *** | PRR vs. MRR | TP 14 | * |
| PRR vs. Sham | TP 14 | *** | PRR vs. Sham | TP 14 | ns | PRR vs. Sham | TP 14 | *** | PRR vs. Sham | TP 14 | *** |
| MRR vs. Sham | TP 14 | *** | MRR vs. Sham | TP 14 | ns | MRR vs. Sham | TP 14 | *** | MRR vs. Sham | TP 14 | *** |
| PRR vs. MRR | TP 21 | ns | PRR vs. MRR | TP 21 | ns | PRR vs. MRR | TP 21 | *** | PRR vs. MRR | TP 21 | ** |
| PRR vs. Sham | TP 21 | *** | PRR vs. Sham | TP 21 | * | PRR vs. Sham | TP 21 | *** | PRR vs. Sham | TP 21 | *** |
| MRR vs. Sham | TP 21 | ns | MRR vs. Sham | TP 21 | ns | MRR vs. Sham | TP 21 | *** | MRR vs. Sham | TP 21 | *** |
| PRR vs. MRR | TP 28 | ** | PRR vs. MRR | TP 28 | ns | PRR vs. MRR | TP 28 | *** | PRR vs. MRR | TP 28 | *** |
| PRR vs. Sham | TP 28 | *** | PRR vs. Sham | TP 28 | ns | PRR vs. Sham | TP 28 | *** | PRR vs. Sham | TP 28 | *** |
| MRR vs. Sham | TP 28 | ns | MRR vs. Sham | TP 28 | ns | MRR vs. Sham | TP 28 | *** | MRR vs. Sham | TP 28 | ns |

**Table S3: Detailed statistics of Figure 2 in the main text**

| | PRR | MRR |
|---|---|---|
| **Best-fit values** | | |
| **Slope** | 0.7864 | 0.9059 |
| **Y-intercept** | -0.9918 | -0.6534 |
| **X-intercept** | 1.261 | 0.7213 |
| **1/slope** | 1.272 | 1.104 |
| **Std. Error** | | |
| **Slope** | 0.04538 | 0.03658 |
| **Y-intercept** | 0.1004 | 0.07162 |
| **95% Confidence Intervals** | | |
| **Slope** | 0.6951 to 0.8778 | 0.8323 to 0.9796 |
| **Y-intercept** | -1.194 to -0.7897 | -0.7977 to -0.5092 |
| **X-intercept** | 1.113 to 1.388 | 0.5995 to 0.8310 |
| **Goodness of Fit** | | |
| **R squared** | 0.8672 | 0.9317 |
| **Sy.x** | 0.2781 | 0.2731 |
| **Is slope significantly non-zero?** | | |
| **F** | 300.4 | 613.4 |
| **DFn, DFd** | 1, 46 | 1, 45 |
| **P value** | <.001 | <.001 |
| **Deviation from zero?** | Significant | Significant |
| **Equation** | Y = 0.7864*X - 0.9918 | Y = 0.9059*X - 0.6534 |
| **Data** | | |
| **Number of X values** | 95 | 47 |
| **Maximum number of Y replicates** | 1 | 1 |
| **Total number of values** | 48 | 47 |
| **Number of missing values** | 47 | 0 |

Are the slopes equal?
F = 4.216. DFn = 1, DFd = 91
P = .043

If the overall slopes were identical, there was a 4.29% chance of randomly choosing data points with slopes this different. You can conclude that the differences between the slopes were significant.

Because the slopes differ so much, it is impossible to test whether the intercepts differ significantly.
